# Targeted deletion of uterine glandular Foxa2 induces embryonic diapause in mice

**DOI:** 10.1101/2022.03.15.484401

**Authors:** Mitsunori Matsuo, Jia Yuan, Yeon Sun Kim, Amanda Dewar, Hidetoshi Fujita, Sudhansu K. Dey, Xiaofei Sun

## Abstract

Embryonic diapause is a reproductive strategy in which embryo development and growth is temporarily arrested within the uterus to ensure the survival of neonates and mothers during unfavorable conditions. Pregnancy is reinitiated when conditions become favorable for neonatal survival. The mechanism of how the uterus enters diapause remains unclear. Mice with uterine deletion of *Foxa2*, a transcription factor, are infertile. In this study, we show that dormant blastocysts are recovered from these mice on day 8 of pregnancy with persistent uterine *Msx1* expression, a gene critical to maintaining the uterine quiescent state, suggesting that these mice enter embryonic diapause. Leukemia inhibitory factor (LIF) can resume implantation in these mice. Although estrogen is critical for implantation in progesterone (P_4_)-primed uterus, our current model reveals that FOXA2-independent estrogenic effects are detrimental to sustaining uterine quiescence. Interestingly, P_4_ and anti-estrogen can prolong uterine quiescence in the absence of FOXA2. While we find that *Msx1* expression persists in the uterus deficient in *Foxa2*, the complex relationship of FOXA2 with *Msx* genes and estrogen receptors remains to be explored.

## Introduction

Embryonic diapause is a reproductive strategy in which embryo development and growth is temporarily arrested within the uterus, but is reinitiated when conditions are favorable for neonatal and maternal survival (1, 2). During diapause, blastocyst growth, DNA synthesis, mitosis, and metabolic activity are temporarily arrested within the uterus, which also becomes quiescent to support blastocyst survival.

The triggers for diapause vary widely across species, ranging from photoperiod, temperature, metabolic stress, lactation, or nutrition (3). It is known to occur in more than 100 species spanning over 7 strata. In mice, experimental ovariectomy on the morning of day 4, before preimplantation estrogen (E) secretion, induces embryonic diapause (4); alternatively, embryonic diapause can be induced by injections of estrogen receptor antagonists on days 3 and 4 of pregnancy to neutralize estrogen function (5). Blastocyst reactivation can be rapidly initiated by a single injection of estrogen in an ovariectomized dormant uterus (4). Preimplantation E secretion on day 4 morning induces Leukemia inhibitory factor (LIF) to initiate implantation. Interestingly, blastocysts in *Lif*^-/-^ females undergo diapause (6). In spite of these recognized factors, the molecular mechanism which initiates embryonic diapause is still not fully understood.

FOXA2 (Forkhead box protein A2), which is expressed in glandular epithelia in the mouse uterus, plays a key role in uterine gland development and implantation. Neonatal deletion of uterine *Foxa2* causes defects in gland development (7). Female mice with uterine glandular deletion of *Foxa2* after puberty have implantation/decidualization failure due to compromised LIF induction on day 4 of pregnancy (8).

In this study, we show that deletion of *Foxa2* in mouse uterine glands causes embryonic diapause. Dormant embryos were retrieved from uteri on day 8 of pregnancy. *Msx1* expression, which appears to be critical to maintain a quiescent uterine environment (5), was maintained in *Foxa2* deficient mice in our studies. Although implantation was triggered by a single LIF injection on day 8 of pregnancy in *Foxa2* deficient mice, these mice are not able to support pregnancy to full term. Furthermore, we found that suppression of estrogenic effects by either progesterone (P_4_) supplement or application of an estrogen receptor antagonist significantly improved the quality of embryonic diapause in *Foxa2* deficient mice. Our study reveals that *Foxa2* plays an important role in mammalian embryonic diapause and that FOXA2-independent E_2_ effects are detrimental to uterine quiescence during diapause.

## Results

### Uterine deletion of Foxa2 results in female infertility due to disrupted Lif induction during implantation

Previous reports have shown that *Foxa2* is expressed in the glandular epithelium before and during pregnancy (7, 9). Uterine glandular *Foxa2* is critical to normal implantation (8). Using mice with uterine specific (*Foxa2^f/f^Pgr*^*Cre*/+^) and uterine epithelial specific deletion of *Foxa2* (*Foxa2^f/f^Ltf*^*Cre*/+^), we confirmed these mice are infertile (Figure 1a) due to the lack of *Lif* induction on day 4 of pregnancy (Figure 1d). Since the timing and domain of Cre activity driven by an *Ltf* or *Pgr* promoter differs (Figure 1b and Supplemental figure 1), *Foxa2^f/f^Ltf*^*Cre*/+^ females maintain FOXA2 expression in uterine glands before puberty (Figure 1c), whereas *Foxa2^f/f^Pgr*^*Cre*/+^ females have a minimal number of glands on postnatal day 30 (Figure 1c) due to early deletion of *Foxa2* (Figure 1b). In mature pregnant females, no positive signals of FOXA2 are observed in either of the two mouse models on day 4 of pregnancy (Figure 1c).

**Figure 1.**
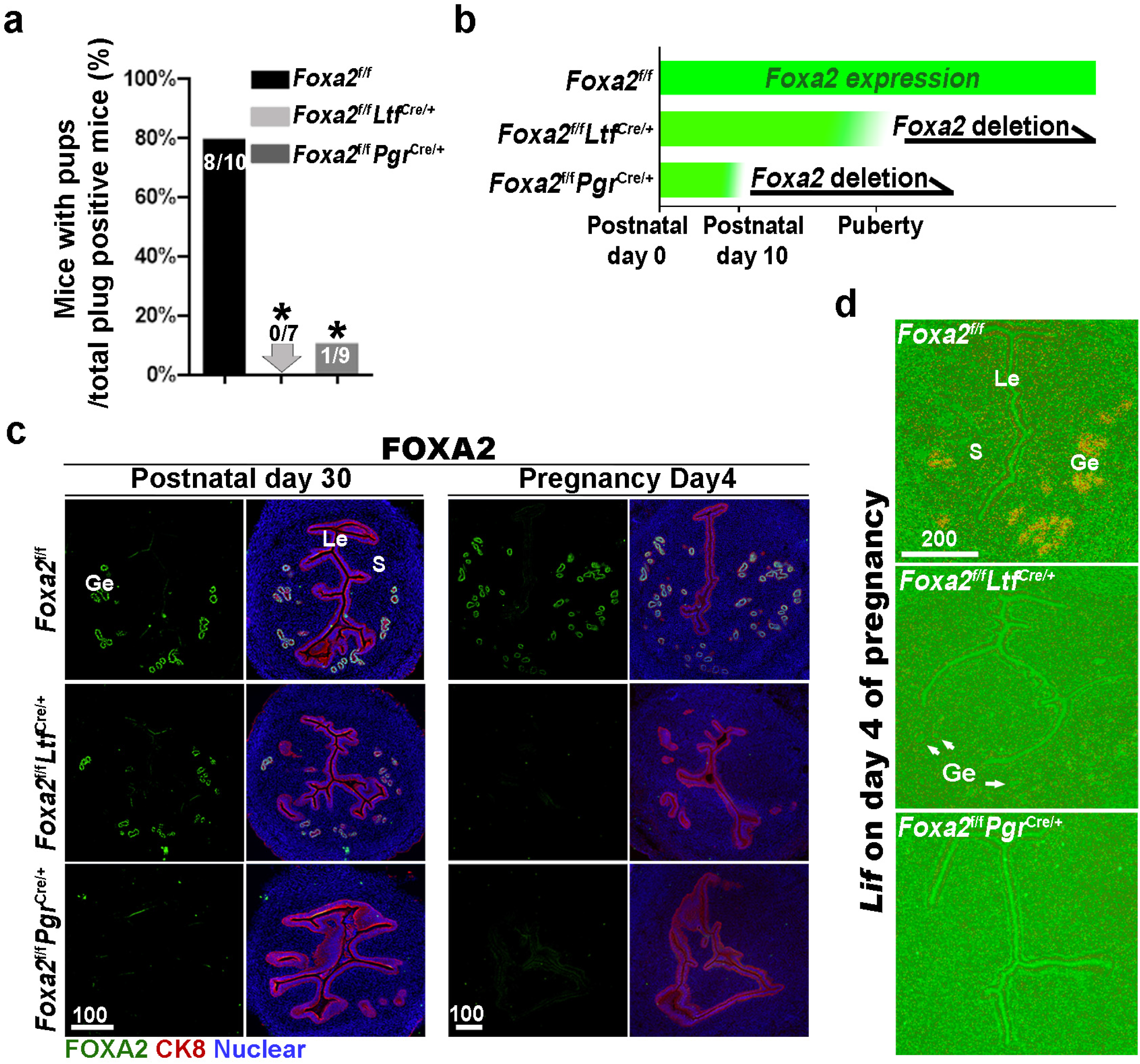
Uterine conditional deletion of *Foxa2* causes loss of LIF secretion and female infertility. a. Percentage of mice with pups per total plug positive mice. Numbers on bars indicate mice with pups over total number of plug positive mice. \**P* < 0.05 by Student’s t-test. b. Cre recombinase activity starts differently in *Foxa2^f/f^Ltf*^*Cre*+^ and *Foxa2^f/f^Pgr*^*Cre*+^ females. c. Immunostaining of FOXA2 in the uteri on postnatal day 30 and day 4 pregnancy of *Foxa2^f/f^, Foxa2^f/f^Ltf^Cre+^* and *Foxa2^f/f^Pgr*^*Cre*+^ females. Epithelial cells are outlined by Cytokeratin 8 (CK8) staining. Scale bars: 100 μm. d. *In situ* hybridization of *Lif* in day 4 of pregnant uteri from *Foxa2^f/f^, Foxa2^f/f^Ltf^Cre+^* and *Foxa2^f/f^Pgr*^*Cre*+^ females. White arrows point to uterine glands. Scale bar: 200 μm. Le, luminal epithelia; Ge, glandular epithelia; S, stroma.

### *Blastocysts enter embryonic diapause in Foxa2^f/f^Ltf*^*Cre*/+^ and *Foxa2^f/f^Pgr*^*Cre*/+^ mice

Suppression of preimplantation estrogen secretion on day 4 of pregnancy renders mouse uteri quiescent to implantation. This uterine status, also called the neutral phase, can be extended by a daily supplement of progesterone. Un-implanted blastocyst development in quiescent uteri is arrested (10–12); in *Lif*^-/-^ females, blastocysts recovered on day 7 of pregnancy retained implantation capabilities once transferred to wild-type surrogate uteri (6). This result suggests that *Lif*^-/-^ uteri are able to maintain the quiescent phase in spite of presumed estrogen secretion on day 4 of pregnancy. Given the absence of LIF induction in *Foxa2* deficient uteri, we next examined whether *Foxa2^f/f^Ltf*^*Cre*/+^ and *Foxa2^f/f^Pgr*^*Cre*/+^ uteri enter diapause after day 4 of pregnancy.

Day 8 uteri were examined for implantation in *Foxa2^f/f^Ltf*^*Cre*/+^, *Foxa2^f/f^Pgr*^*Cre*/+^ and control (*Foxa2^f/f^*) females. *Foxa2^f/f^* uteri show implantation sites with apparently normal morphology (Figure 2a). In contrast, implantation sites were rarely observed in *Foxa2^f/f^Ltf*^*Cre*/+^ or *Foxa2^f/f^Pgr*^*Cre*/+^ females (Figure 2a). The rate of plug-positive females with implantation sites in *Foxa2^f/f^Ltf*^*Cre*/+^ and *Foxa2^f/f^Pgr*^*Cre*/+^ mice was 16.7% (1/6) and 42.9% (3/7), respectively, significantly lower than the rate of *Foxa2^f/f^* mice (100%, 6/6) (Supplemental Table1).

**Figure 2.**
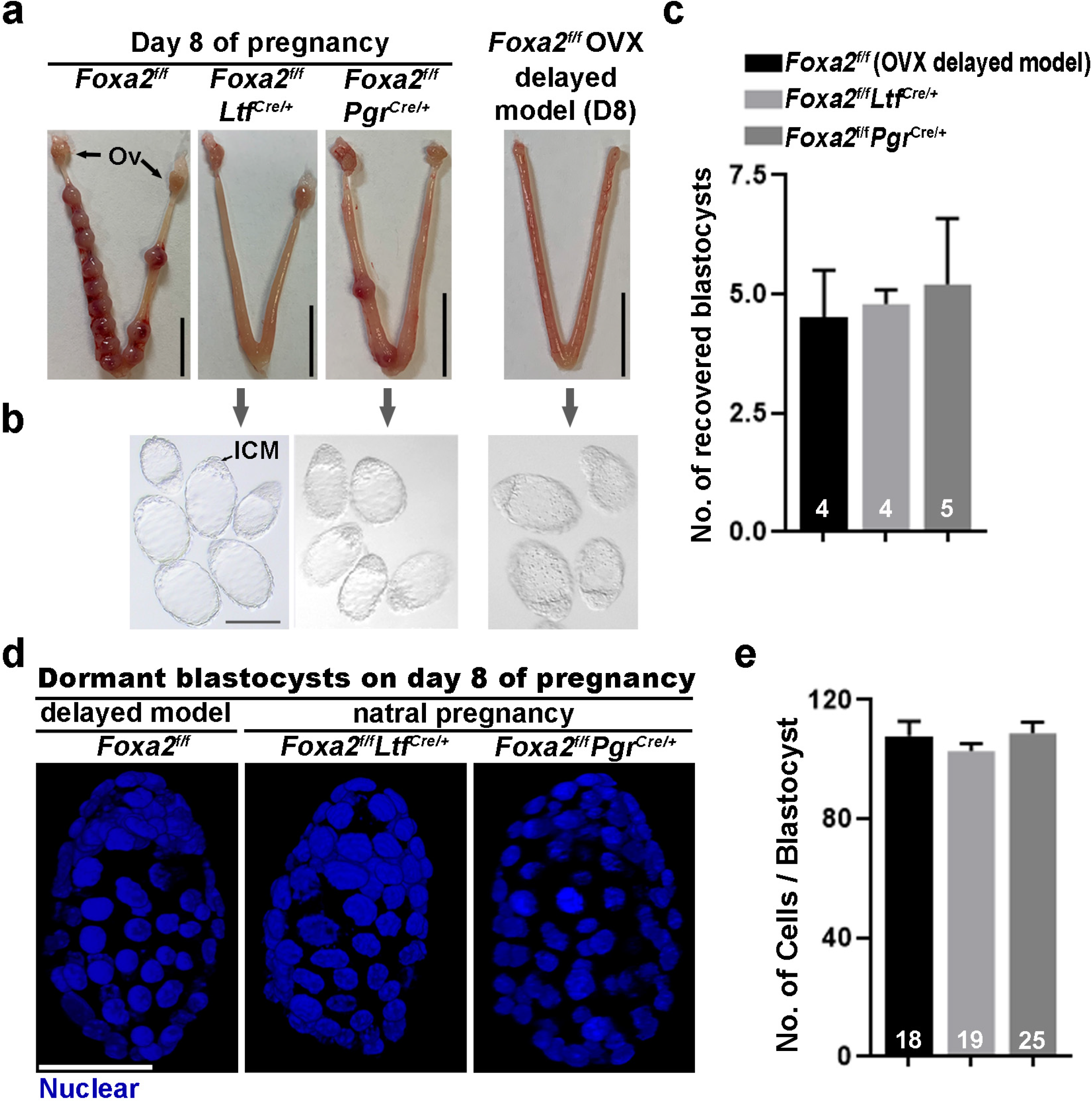
Dormant blastocysts are present in *Foxa2^f/f^Ltf*^*Cre*+^ and *Foxa2^f/f^Pgr*^*Cre*+^ uteri on day 8 pregnancy. a. Representative photographs of day 8 pregnant uteri from *Foxa2^f/f^, Foxa2^f/f^Ltf^Cre+^* and *Foxa2^f/f^Pgr*^*Cre*+^ females. An ovariectomy-induced delayed model of *Foxa2^f/f^* mice served as prototypical control in maintaining dormant blastocysts. Scale bar, 10 mm. Ov, ovary. b. Blastocysts recovered from *Foxa2^f/f^Ltf*^*Cre*+^ and *Foxa2^f/f^Pgr*^*Cre*+^ uteri on day 8. Blastocysts retrieved from ovariectomized *Foxa2^f/f^* mice in delay served as controls. ICM, inner cell mass. Scale bar, 100 μm. Quantification of blastocyst numbers were shown in penal c. Numbers on bars indicate numbers of animals examined. Values are expressed as mean + SEM. d. Representative photographs of nuclear staining of dormant blastocysts recovered from mice without implantation sites. Scale bar, 50 μm. e. Average cell numbers per blastocyst. Numbers of embryos examined are shown on bars. Values are expressed as mean + SEM.

Dormant blastocysts were recovered by flushing *Foxa2^f/f^Ltf*^*Cre*/+^ and *Foxa2^f/f^Pgr*^*Cre*/+^ uterine horns on day 8 of pregnancy (Figure 2b). A diapause model created by ovariectomy on day 4 of pregnancy was used as a prototypical control (Supplemental figure 2), in which ~4.5 dormant blastocysts were retrieved. A similar number of blastocysts is recovered from *Foxa2^f/f^Ltf*^*Cre*/+^ and *Foxa2^f/f^Pgr*^*Cre*/+^ uterine horns on day 8 of pregnancy (Figure 2c), and they morphologically resembled the dormant blastocysts from the ovariectomy delayed model (Figure 2b). It is known that dormant blastocysts cease mitotic activity and cell proliferation (5). Using DAPI staining (Figure 2d), we found that blastocysts retrieved from *Foxa2^f/f^Ltf*^*Cre*/+^ and *Foxa2^f/f^Pgr*^*Cre*/+^ uteri have comparable cell numbers to those recovered from diapausing uteri achieved by ovariectomy (Figure 2e).

### *Foxa2^f/f^Ltf*^*Cre*/+^ and *Foxa2^f/f^Pgr*^*Cre*/+^ uteri show characteristics of uterine quiescence

Uterine quiescence apparently depends on the presence of muscle segment homeobox (*Msx*) genes. In mice and other diapausing animals, *Msx1* and *Msx2* genes persist during diapause, but their levels are quickly suppressed with blastocyst reactivation and implantation (5). However, mice with uterine conditional deletion of both *Msx1* and *Msx2* fail to achieve diapause and reactivation (5, 13). Since dormant blastocysts were recovered from *Foxa2^f/f^Ltf*^*Cre*/+^ and *Foxa2^f/f^Pgr*^*Cre*/+^ uteri on day 8 of pregnancy, we suspected that *Foxa2^f/f^Ltf*^*Cre*/+^ and *Foxa2^f/f^Pgr*^*Cre*/+^ uteri remain quiescent in the absence of LIF induction. We examined *Msx1* expression in the uterus on days 4 and 8 of pregnancy by fluorescence *in situ* hybridization. *Msx1* signals were observed in epithelial cells before the estrogen surge on day 4 in *Foxa2^f/f^* uteri; whereas luminal epithelial *Msx1* signals were suppressed after the estrogen secretion (Figure 3a) (14). Remarkably, *Msx1* expression persisted in *Foxa2^f/f^Ltf*^*Cre*/+^ and *Foxa2^f/f^Pgr*^*Cre*/+^ luminal epithelial cells on day 8 of pregnancy (Figure 3a). Epidermal growth factor receptor (EGFR) is present in day 4 blastocysts, but becomes suppressed during dormancy (15). This is consistent with our current findings that EGFR expression is significantly lower in blastocysts recovered from *Foxa2^f/f^Ltf*^*Cre*/+^ and *Foxa2^f/f^Pgr*^*Cre*/+^ uteri on day 8 of pregnancy as compared to those retrieved from *Foxa2^f/f^* uteri in the evening of day 4 (Figure 3b). The mitotic activity in trophectoderm of the recovered blastocysts in *Foxa2^f/f^Ltf*^*Cre*/+^ and *Foxa2^f/f^Pgr*^*Cre*/+^ mice on day 8 is also arrested (Figure 3c). These results suggest *Foxa2^f/f^Ltf*^*Cre*/+^ and *Foxa2^f/f^Pgr*^*Cre*/+^ uteri remain quiescent until at least day 8 of pregnancy, providing a uterine environment suitable for embryonic diapause.

**Figure 3.**
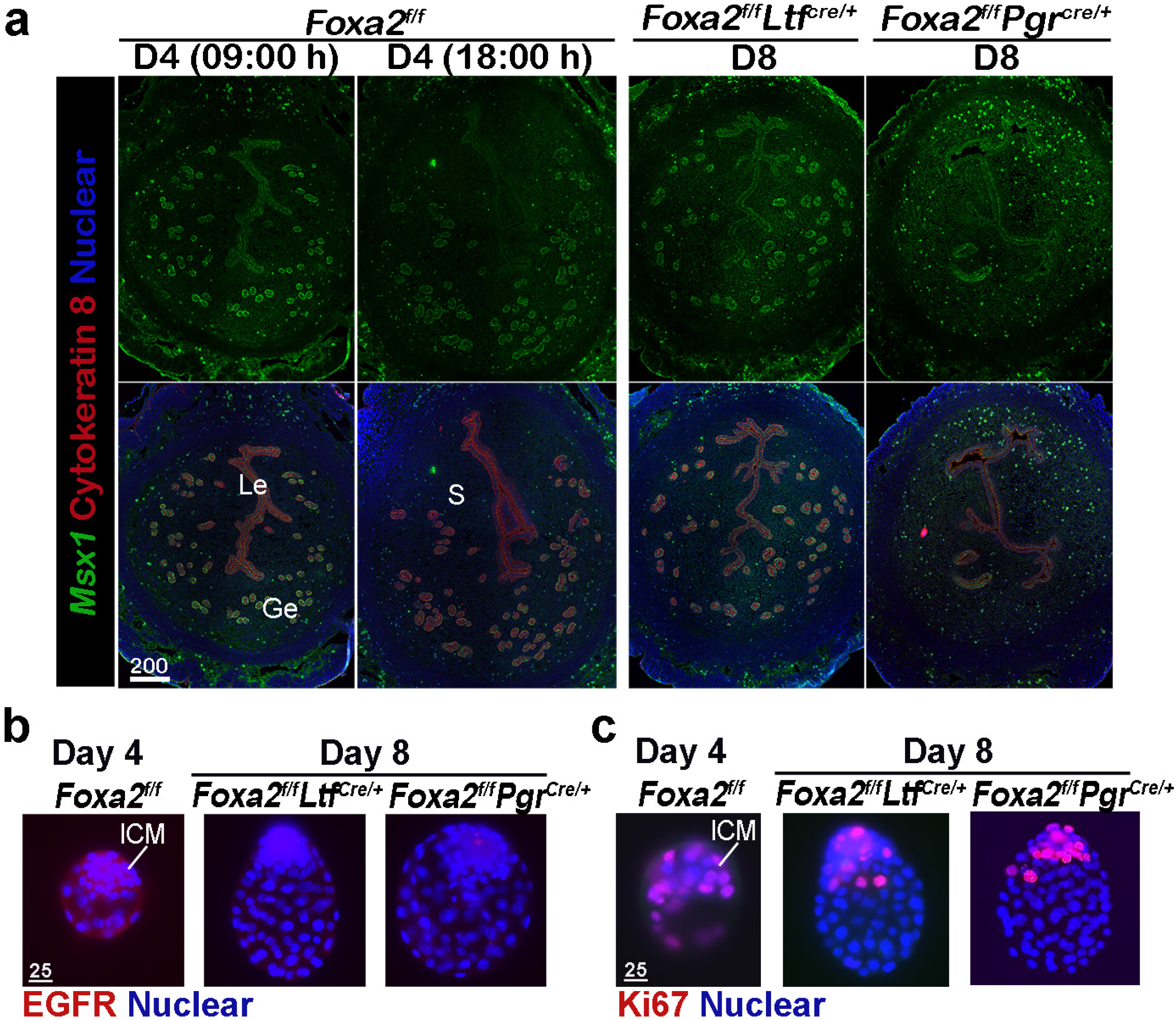
*Foxa2^f/f^Ltf*^*Cre*+^ and *Foxa2^f/f^Pgr*^*Cre*+^ females maintain uterine quiescence when examined on day 8 of pregnancy. a. Fluorescence *in situ* hybridization of *Msx1* in days 4 and 8 pregnant uteri from *Foxa2^f/f^, Foxa2^f/f^Ltf^Cre+^* and *Foxa2^f/f^Pgr*^*Cre*+^ females. Scale bar: 200 μm. b. EGFR immunostaining on dormant blastocysts. Positive signals were observed in activated blastocysts recovered from *Foxa2^f/f^* uteri on day 4 of pregnancy. Scale bar: 25 μm. c. Ki67 immunostaining on dormant blastocysts collected from day 8 *Foxa2^f/f^Ltf*^*Cre*+^ and *Foxa2^f/f^Pgr*^*Cre*+^ females. Scale bar: 25 μm. ICM, inner cell mass.

### *Foxa2^f/f^Ltf*^*Cre*/+^ and *Foxa2^f/f^Pgr^Cre/+^ uterine readiness to be activated deteriorates during diapause*

Embryonic diapause and uterine quiescence are reversible with a single injection of estrogen or LIF in mice (4, 16). Implantation failure has been shown to be rescued in *Foxa2^f/f^Ltf*^*Cre*/+^ and *Foxa2^f/f^Pgr*^*Cre*/+^ females by LIF administration on day 4 (8). No further analysis was carried out. Since dormant blastocysts are recovered from *Foxa2^f/f^Ltf*^*Cre*/+^ and *Foxa2^f/f^Pgr*^*Cre*/+^ uteri on day 8 of pregnancy in our studies, we examined if the diapausing blastocysts can be rejuvenated. *Foxa2^f/f^Ltf*^*Cre*/+^ and *Foxa2^f/f^Pgr*^*Cre*/+^ females received one injection of recombinant LIF (20 μg/mouse) on day 4 or 8, and the uteri were examined two days later (Figure 4a). Consistent with the previous report (8), implantation sites were observed in both *Foxa2^f/f^Ltf*^*Cre*/+^ and *Foxa2^f/f^Pgr*^*Cre*/+^ uteri if LIF was given on day 4 of pregnancy (Figure 4b). Further histological evaluations revealed that normal-looking implantation chambers formed similar to those in *Foxa2^f/f^* mice on day 6 of pregnancy (Figures 4b and c). Almost all *Foxa2^f/f^Ltf*^*Cre*/+^ females (5 of 6) possessed implantation sites, whereas less than half (4 of 9) *Foxa2^f/f^Pgr*^*Cre*/+^ females had implantation sites (Figure 4d), although the number of implantation sites were comparable in pregnant females.

**Figure 4.**
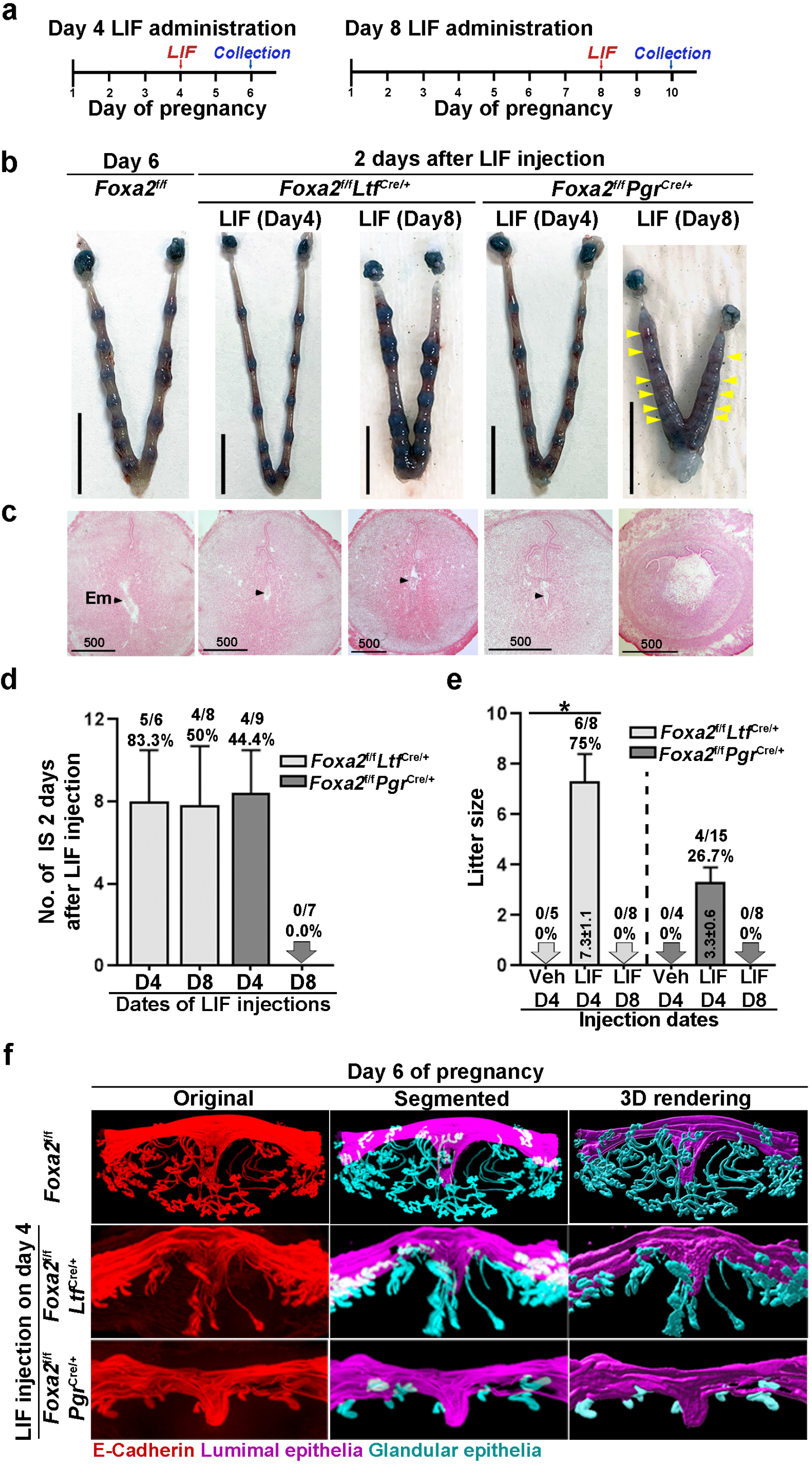
Pregnancy in *Foxa2^f/f^Ltf*^*Cre*+^ and *Foxa2^f/f^Pgr*^*Cre*+^ females with LIF treatment. a. Schematic outline of sample collection. LIF, LIF administration (20 μg). b. Representative photograph of uteri from *Foxa2^f/f^Ltf*^*Cre*+^ and *Foxa2^f/f^Pgr*^*Cre*+^ females (days 6 and 10) with LIF treatment. *Foxa2^f/f^* uteri on day 6 serve as control. Scale bar, 10 mm. Histological pictures of implantation sites in panel b were presented in panel c. Arrowheads point to embryos. Em, embryo. Scale bar, 500 μm. d. Average number of implantation sites in *Foxa2^f/f^Ltf*^*Cre*+^ and *Foxa2^f/f^Pgr*^*Cre*+^ mice treated with LIF (20 μg) on day 4 or 8 of pregnancy. Numbers and percentage on bars indicate mice with implantation sites over total number of plug positive mice. e. Litter sizes of *Foxa2^f/f^Ltf*^*Cre*+^ and *Foxa2^f/f^Pgr*^*Cre*+^ mice treated with LIF (20 μg) or vehicle on days 4 or 8 of pregnancy. Numbers and percentage on bars indicate mice with pups over total number of plug positive mice. \**P*<0.05. f. 3D visualization of day 6 implantation sites in *Foxa2^f/f^, Foxa2^f/f^Ltf^Cre+^* and *Foxa2^f/f^Pgr*^*Cre*+^ females. Images of E-cadherin immunostaining, segmented, and 3D rendered images of day 6 implantation sites in each genotype show defects in *Foxa2^f/f^Pgr*^*Cre*+^ females with a LIF injection on day 4 of pregnancy.

Notably, implantation sites with a normal appearance were observed in *Foxa2^f/f^Ltf*^*Cre*/+^ uteri when LIF was given on day 8 of pregnancy (Figure 4b), albeit only faint blue bands were seen in *Foxa2^f/f^Pgr*^*Cre*/+^ uteri. Histology of implantation sites confirmed this observation. Implantation chambers form in *Foxa2^f/f^Ltf*^*Cre*/+^ implantation sites, but neither embryos nor implantation chambers were found in *Foxa2^f/f^Pgr*^*Cre*/+^ implantation sites (Figure 4c). The rate of *Foxa2^f/f^Ltf*^*Cre*/+^ females with implantation sites decreased from 83.3% to 50% in females receiving LIF injection on day 4 (Figure 4d). All *Foxa2^f/f^Pgr*^*Cre*/+^ females showed abnormal light blue bands without recognizable implantation chambers when examined two days after LIF injection; the implantation chambers showed no further development (Figure 4d).

Glands have been shown to be essential for implantation and pregnancy success (8, 17). FOXA2 plays a key role in mouse uterine glandular genesis, and neonatal deletion of *Foxa2* in mouse uteri causes defects in gland development (7). To examine glands, day 6 implantation sites of *Foxa2^f/f^Ltf*^*Cre*/+^ and *Foxa2^f/f^Pgr*^*Cre*/+^ females with LIF injection on day 4 were stained with an E-Cadherin antibody. Tridimensional images were acquired as previously described (18). The number of glands in *Foxa2^f/f^Ltf*^*Cre*/+^ implantation sites was significantly reduced compared to those in natural day 6 implantation sites of *Foxa2^f/f^* females (Figure 4f). As previously reported (7), glands were rarely observed in *Foxa2^f/f^Pgr*^*Cre*/+^ implantation sites. These data suggest that an increased number of glands is not required for uterine quiescence and embryonic diapause, but the presence of a minimal number of glands is critical for reactivation after diapause.

In mammalian embryonic diapause, arrest of blastocyst development and uterine quiescence are transitory. Upon reactivation, the uterine environment is sufficient to support embryo development to term when conditions are favorable for neonatal survival. To study whether reactivated uteri in *Foxa2^f/f^Ltf*^*Cre*/+^ and *Foxa2^f/f^Pgr*^*Cre*/+^ females are able to support full-term pregnancy, litter sizes were counted. Six of eight *Foxa2^f/f^Ltf^Cre/+^*, but only 4 of 15 *Foxa2^f/f^Pgr*^*Cre*/+^ females injected with LIF on day 4 successfully delivered live pups, and *Foxa2^f/f^Pgr*^*Cre*/+^ females have reduced litter sizes (Figure 4e). Notably, neither *Foxa2^f/f^Ltf*^*Cre*/+^ nor *Foxa2^f/f^Pgr*^*Cre*/+^ females with day 8 LIF injection were able to support full-term pregnancy, in spite of implantation occurring two days after LIF injection in *Foxa2^f/f^Ltf*^*Cre*/+^ females (Figure 4e). These results suggest that uterine readiness for reactivation in *Foxa2^f/f^Ltf*^*Cre*/+^ and *Foxa2^f/f^Pgr*^*Cre*/+^ deteriorates during diapause with preimplantation estrogen secretion.

### Progesterone supplement during diapause improves pregnancy outcomes in Foxa2^f/f^Ltf^Cre/+^ and Foxa2^f/f^Pgr^Cre/+^ females after reactivation

Progesterone is required to maintain uterine quiescence and blastocyst viability in mouse embryonic diapause. Embryonic diapause is also experimentally induced in the mouse by ovariectomy on day 4 of pregnancy before estrogen secretion and maintained by daily P_4_ injections (4, 19). To examine if P_4_ levels decrease without implantation in *Foxa2^f/f^Ltf*^*Cre*/+^ and *Foxa2^f/f^Pgr*^*Cre*/+^ females, we evaluated serum concentration of P_4_ on days 4 and 8 of pregnancy and estrogen on day 4 pregnancy in *Foxa2^f/f^Ltf^Cre/+^, Foxa2^f/f^Pgr*^*Cre*/+^ and *Foxa2^f/f^* females. P_4_ and E_2_ levels in *Foxa2^f/f^Ltf*^*Cre*/+^ and *Foxa2^f/f^Pgr*^*Cre*/+^ females were comparable to those in *Foxa2^f/f^* mice (Supplemental figures 3a and b).

Although *Foxa2^f/f^Ltf*^*Cre*/+^ and *Foxa2^f/f^Pgr*^*Cre*/+^ females have normal P_4_ and E_2_ levels, the uterine edema in *Foxa2^f/f^Pgr*^*Cre*/+^ females two days after LIF injection on day 8 (Figure 4b) suggests augmented estrogenic effects during diapause. Therefore, we administered P_4_ on days 5, 7 and 9 with LIF injection on day 8 to counter the increased estrogenic effects in *Foxa2^f/f^Ltf*^*Cre*/+^ and *Foxa2^f/f^Pgr*^*Cre*/+^ females (Figure 5a). Embryo Implantation was evaluated two days after LIF administration. All *Foxa2^f/f^Ltf*^*Cre*/+^ and *Foxa2^f/f^Pgr*^*Cre*/+^ females had implantation sites with distinct blue bands (Figures 5b and d). Histological analysis identified embryos in the implantation chambers in mice of both genotypes (Figure 5c). Implantation rates and the numbers of implantation sites appear normal (Figure 5d). A comparable decidual response as revealed by RNA levels of *Bmp2* is observed in these *Foxa2^f/f^Ltf*^*Cre*/+^ females in day 6 *Foxa2^f/f^* implantation sites (Supplemental figures 4). Furthermore, around 40% of *Foxa2^f/f^Ltf*^*Cre*/+^ and *Foxa2^f/f^Pgr*^*Cre*/+^ females successfully delivered progeny, although litter sizes were small (2~3 pups/litter) (Figure 5e). These data suggest P_4_ supplementation improves uterine conditions during *Foxa2^f/f^Ltf*^*Cre*/+^ and *Foxa2^f/f^Pgr*^*Cre*/+^ diapause.

**Figure 5.**
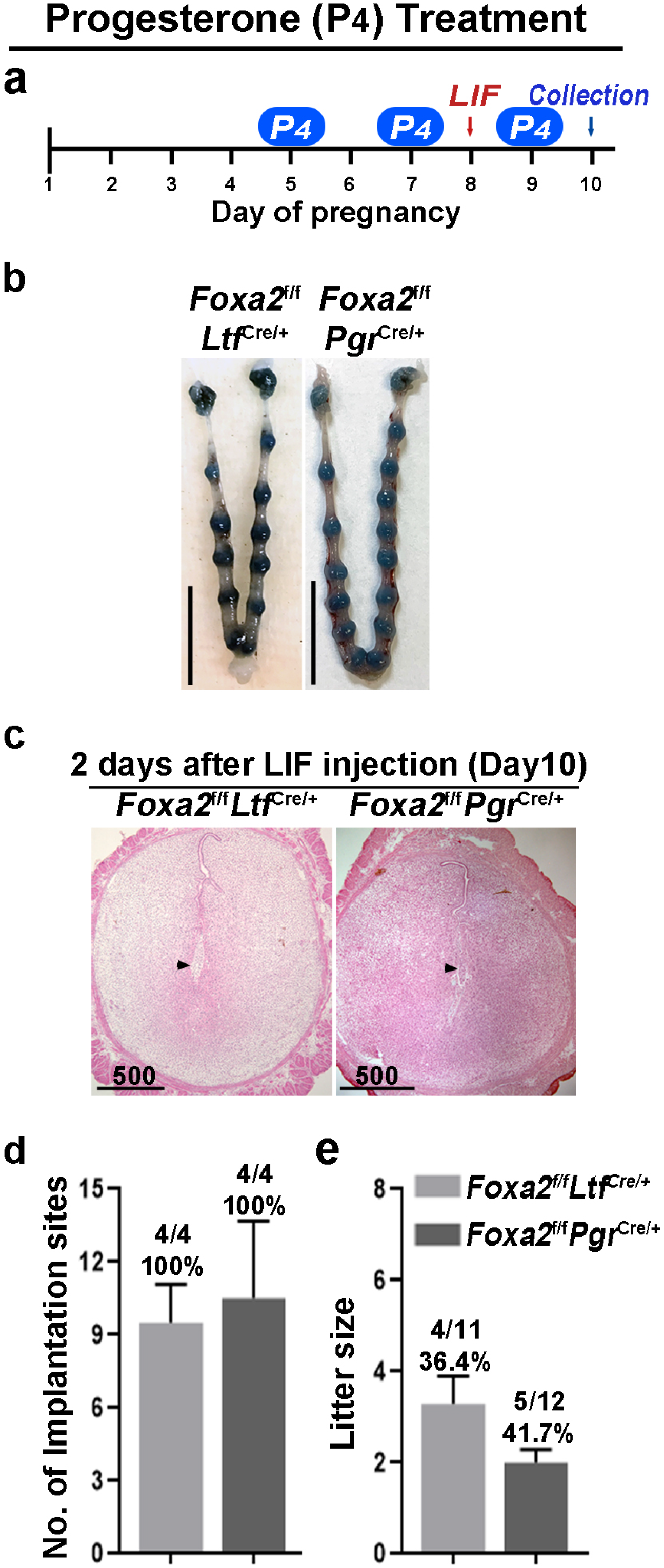
Counterbalance of estrogenic effects by P_4_ improves maintenance of diapause in *Foxa2^f/f^Ltf*^*Cre*+^ and *Foxa2^f/f^Pgr*^*Cre*+^ mice. a. Scheme of P_4_ treatment. *Foxa2^f/f^Ltf*^*Cre*+^ and *Foxa2^f/f^Pgr*^*Cre*+^ mice were treated with LIF (20 μg) on day 8. Pregnancy was evaluated on day 10, two days after LIF administration. b. Representative photographs of uteri in *Foxa2^f/f^Ltf*^*Cre*+^ and *Foxa2^f/f^Pgr*^*Cre*+^ mice with P_4_ supplement two days after LIF administration. Scale bar, 10 mm. Histological pictures of implantation sites in panel b were presented in panel c. Scale bar, 500 μm. d. Average number of implantation sites in *Foxa2^f/f^Ltf*^*Cre*+^ and *Foxa2^f/f^Pgr*^*Cre*+^ mice with P_4_ supplement. Numbers and percentage on bars indicate mice with implantation sites over total number of plug positive mice. e. Litter sizes of *Foxa2^f/f^Ltf*^*Cre*+^ and *Foxa2^f/f^Pgr*^*Cre*+^ mice with P_4_ supplement. Numbers and percentage on bars indicate mice with pups over total number of plug positive mice.

### Suppression of estrogen action during diapause improves pregnancy outcomes in Foxa2^f/f^Ltf^Cre/+^ and Foxa2^f/f^Pgr^Cre/+^ females after reactivation

In mice, preimplantation estrogen secretion on day 4 of pregnancy triggers a transitory receptive phase in uteri; if implantation does not occur, the receptive phase shifts to a refractory phase (20). This activity suggests that estrogen has a biphasic effect on embryo implantation: a positive effect to induce uterine receptivity and a negative effect in changing the receptive uterus to a non-receptive state. In *Foxa2^f/f^Ltf*^*Cre*/+^ and *Foxa2^f/f^Pgr*^*Cre*/+^ females, although estrogen induced LIF expression was abolished, FOXA2 independent negative estrogenic effects may gradually induce the refractory phase in uteri. To test this possibility, we administrated an estrogen receptor antagonist (ICI-182,780, named ICI) on days 3, 5 and 7 before day 8 LIF injection in *Foxa2^f/f^Ltf*^*Cre*/+^ and *Foxa2^f/f^Pgr*^*Cre*/+^ females (Figure 6a). Of note, embryonic diapause can also be experimentally induced in mice via 50 μg ICI injections on days 3 and 4 (5). We have also confirmed this observation in our present study. To avoid the suppression of E_2_-induced LIF secretion on day 4, the dose of ICI was lowered to 25 μg per injection, the level at which implantation occurs normally in *Foxa2^f/f^* females (Figure 6b).

**Figure 6.**
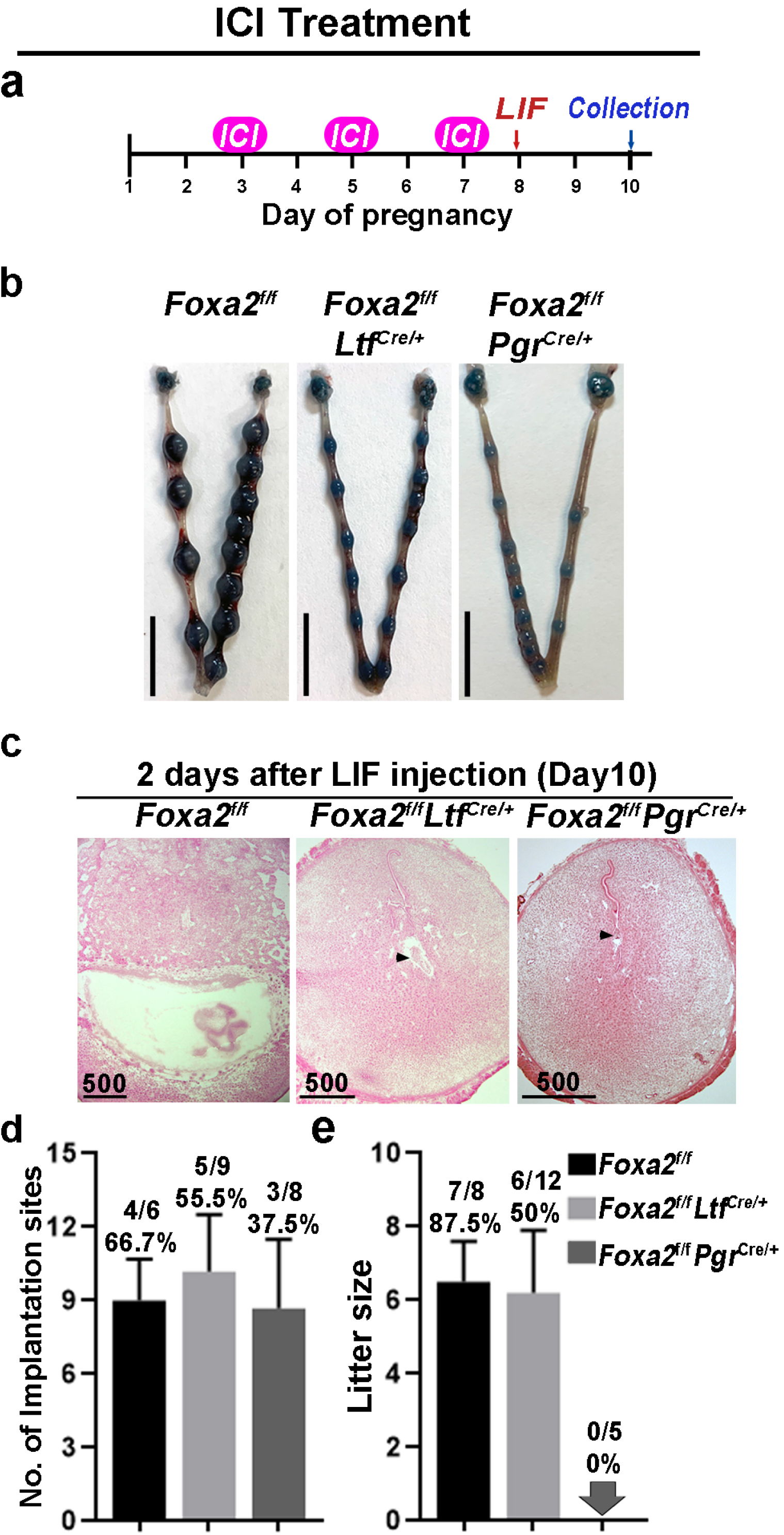
Counterbalance of estrogenic effects by ICI improves diapause in *Foxa2^f/f^Ltf*^*Cre*+^ and *Foxa2^f/f^Pgr*^*Cre*+^ mice. a. Scheme of of ICI treatment. *Foxa2^f/f^Ltf*^*Cre*+^ and *Foxa2^f/f^Pgŕ^Cre+^* mice were treated with LIF (20 μg) on day 8. Pregnancy was evaluated on day 10, two days after LIF administration. b. Representative photographs of uteri in *Foxa2^f/f^Ltf*^*Cre*+^ and *Foxa2^f/f^Pgr*^*Cre*+^ mice with ICI treatment two days after LIF administration. *Foxa2^f/f^* mice have normal day 10 implantation sites, suggesting implantation occurs under 25 μg ICI treatment in *Foxa2^f/f^* mice. Scale bar, 10 mm. Histological pictures of implantation sites in panel b were presented in panel c. Scale bar, 500 μm. d. Average number of implantation sites in *Foxa2^f/f^Ltf*^*Cre*+^ and *Foxa2^f/f^Pgr*^*Cre*+^ mice with ICI treatment. Numbers and percentage on bars indicate mice with implantation sites over total number of plug positive mice. e. Litter sizes of *Foxa2^f/f^Ltf*^*Cre*+^ and *Foxa2^f/f^Pgr*^*Cre*+^ mice with ICI treatment. Numbers and percentage on bars indicate mice with pups over total number of plug positive mice.

Similar to P_4_ supplement, ICI treatment improved uterine responses to LIF-induced reactivation of embryos in *Foxa2^f/f^Ltf*^*Cre*/+^ and *Foxa2^f/f^Pgr*^*Cre*/+^ females. Implantation sites with distinct blue bands were observed in *Foxa2^f/f^Ltf*^*Cre*/+^ and *Foxa2^f/f^Pgr*^*Cre*/+^ females two days after LIF injection (Figures 6b). Embryos were identified in implantation chambers in both *Foxa2^f/f^Ltf*^*Cre*/+^ and *Foxa2^f/f^Pgr*^*Cre*/+^ mice (Figure 6c). Quantitatively, 55.5% of *Foxa2^f/f^Ltf*^*Cre*/+^ females and 37.5% of *Foxa2^f/f^Pgr*^*Cre*/+^ females had implantation sites; the number of implantation sites in these mice is comparable to those in *Foxa2^f/f^* females (Figure 6d). A comparable decidual response as indicated by *Bmp2* expression is observed in ICI treated *Foxa2^f/f^Ltf*^*Cre*/+^ females in day 6 *Foxa2^f/f^* implantation sites (Supplemental figures 4). Surprisingly, 50% of *Foxa2^f/f^Ltf*^*Cre*/+^ females supported pregnancy to full-term with a litter size comparable to those of *Foxa2^f/f^* females (Figure 6e). However, no delivery was observed in *Foxa2^f/f^Pgr*^*Cre*/+^ females. These results suggest that a low level of ICI suppressed adverse estrogenic effects on uterine quiescence during diapause.

## Discussion

Over 130 mammalian species experience diapause. The triggers for diapause across species vary widely including sucking stimuli, photoperiod, the availability of nutrition and so forth (11). The uterus is perhaps a determining factor for embryonic diapause in that a non-diapausing embryo undergoes dormancy in a diapausing uterus (21). During diapause, mammals temporarily arrest blastocyst development and metabolic activity within the uterus. In normal pregnancy, uterine sensitivity to implantation is classified into three phases in mice: prereceptive, receptive and nonreceptive (refractory) (22). Mouse uteri attain quiescence in diapause directly from the prereceptive phase via suppression of preimplantation E_2_ secretion or LIF on day 4 of pregnancy (4, 6, 23, 24). However, the mechanism to induce embryonic diapause in mice is not clearly understood. In the current study, we show that suppression of uterine *Foxa2* triggers the mouse uterus to enter a quiescent status, which supports the arrest of embryonic development. A previous study showed that *Foxa2^f/f^Ltf*^*Cre*/+^ females have implantation failure due to LIF suppression prior to implantation (8), which potentially explains why *Foxa2^f/f^Ltf*^*Cre*/+^ uteri are quiescent, since a single injection of LIF is sufficient to initiate embryo implantation. Dormant blastocysts were recovered from *Lif*^-/-^ females on day 7 of pregnancy (6).

Uterine deletion of *Foxa2* in either *Foxa2^f/f^Ltf*^*Cre*/+^ or *Foxa2^f/f^Pgr*^*Cre*/+^ females is not sufficient to maintain complete uterine quiescence by suppressing all uterine metabolic activities. In diapause, quiescent uteri are readily reactivated by an injection of E_2_ or LIF. However, our current study showed that only *Foxa2^f/f^Ltf*^*Cre*/+^ and *Foxa2^f/f^Pgr*^*Cre*/+^ females with day 4 LIF injection delivered progeny; mice with day 8 LIF injection were unable to carry to term. Although *Msx1* persists in *Foxa2^f/f^Ltf*^*Cre*/+^ or *Foxa2^f/f^Pgr*^*Cre*/+^ uteri on day 8 of pregnancy, mice with day 8 LIF injections failed to continue pregnancy to full term, indicating the reactivation capacity for uterine readiness has been deteriorating from day 4 to day 8. Compared with ovariectomy-induced diapause, *Foxa2^f/f^Ltf*^*Cre*/+^ and *Foxa2^f/f^Pgr*^*Cre*/+^ females still have estrogen secretion on day 4 morning. Although E_2_-induced LIF expression is diminished, it is possible that E_2_ continues to have some effects, independent of FOXA2 that slowly compromises uterine readiness for reactivation in *Foxa2^f/f^Ltf*^*Cre*/+^ and *Foxa2^f/f^Pgr*^*Cre*/+^ females. Furthermore, we show that detrimental estrogenic effects on diapause could be countered by P_4_ supplement or ICI treatment.

The exact role of *Foxa2* in LIF induction by E_2_ is not clearly understood. FOXA2 is a nuclear transcription factor and is involved in cell commitment, differentiation, and gene transcription in various organs including the lung, liver, pancreas, and gastrointestinal tract (25, 26). Since both FOXA2 and estrogen receptors (ERs) are transcription factors, it is possible that FOXA2 and ER synergistically turn on LIF activity. On the other hand, ER is activated by estrogen secretion in the morning of day 4, whereas *Foxa2* is constantly expressed in uterine glands, suggesting an alternative possibility that FOXA2 primes transcriptional regulatory regions of *Lif*, thus enabling ER binding. In fact, previous reports showed that FOXA2 is required for chromatin opening during endoderm differentiation (27) and for proper chromatin remodeling in human pancreas specification (28).

Estrogen is harmful to uterine quiescence in mouse diapause. Estrogen is critical for the transition from the prereceptive to the receptive phase in P_4_ primed uteri (22). Without implantation, mouse uteri enter the refractory phase after a short receptive phase (implantation window), suggesting that estrogen quickly terminates the uterine receptive phase. There is evidence that estrogen concentration determines the duration of the uterine receptive phase, wherein a high dose of estrogen shortens the receptive period (29). Conversely, mouse uteri in diapause maintain activation readiness by a bolus shot of estrogen as in the prereceptive phase. Ovariectomy-induced embryonic diapause could last for weeks in mice with continued P_4_ treatment (29, 30), suggesting uterine quiescence can be maintained for significant periods of time in the absence of E_2_. In *Foxa2^f/f^Ltf*^*Cre*/+^ and *Foxa2^f/f^Pgr*^*Cre*/+^ females, LIF induction is suppressed, which avoids a quick switch to the refractory phase. However, the FOXA2 independent estrogen effect remains in *Foxa2^f/f^Ltf*^*Cre*/+^ and *Foxa2^f/f^Pgr*^*Cre*/+^ uteri, slowly compromising arrest of embryonic development and uterine quiescence. This dysfunction is further supported by our finding that P_4_ or a low dose of ICI, which suppress estrogen function, improves the diapause condition in *Foxa2^f/f^Ltf*^*Cre*/+^ and *Foxa2^f/f^Pgr*^*Cre*/+^ females. These results indicate that estrogenic effects are not conducive to mouse diapause.

## Materials and Methods

### Animal and treatment

*Foxa2^f/f^* mice on a CD1 background were generated as described (31). *Foxa2^f/f^Ltf*^*Cre*+^ and *Foxa2^f/f^Pgr^Cre+^* mice were generated by mating *Foxa2^f/f^* females with *Ltf^Cre/+^* males (C57BL/6 and albino B6 mixed background) and *Pgr^Cre/+^* mice. *Ltf*^Cre/+^ and *Pgr^Cre/+^* mice on a C57BL/6 background were generated as described (32, 33). *Foxa2^f/f^, Foxa2^f/f^Ltf^Cre+^* and *Foxa2^f/f^Pgr^Cre+^* mice were housed in the animal care facility at Cincinnati Children’s Hospital Medical Center according to the National Institute of Health and institutional guidelines for laboratory animals. All protocols were approved by the Cincinnati Children’s Animal Care and Use Committee. Mice were provided with autoclaved Laboratory Rodent Diet 5010 (Purina) and UV light-sterilized reverse osmosis/deionized constant-circulation water ad libitum. All mice used in this study were housed under a 12:12 hour light:dark cycle. At least three mice from each genotype were used for each individual experiment.

### Analysis of pregnancy events

Three adult (3-month old) females from each genotype were randomly chosen and housed with a *Foxa2^f/f^* fertile male overnight in separate cages; the morning of finding the presence of a vaginal plug was considered successful mating (day 1 of pregnancy). Plug-positive females were selected for pregnancy experiments. For analysis of parturition, parturition events were monitored from day 18 through day 27 by observing mice daily, morning, noon, and evening.

Litter size, pregnancy rate, gestation length and outcomes were monitored. Implantation sites were examined on pregnancy day 6 or day 8. Blue reaction was performed by intravenous injection of a blue dye solution (Chicago Blue dye) 4 min before mice were sacrificed. Distinct blue bands along the uterus indicated implantation sites. For confirmation of pregnancy in mice showing no blue bands, one uterine horn was flushed with saline for the presence of blastocysts. If blastocysts were present, the contralateral horn was used for experiments; mice without any blastocysts were excluded. ICI (Fulvestrant, Sigma-Aldrich, 25 μg/mouse/day) or progesterone (P_4_, Sigma-Aldrich, 2mg/mouse/day) was administered in the morning (0900h). To induce implantation, a single injection of recombinant leukemia inhibitory factor (LIF, 20μg per mouse) was administrated in the morning (0900h). Embryo implantation sites were examined two days after LIF injection by intravenous injection of a blue dye solution.

### Histology

Tissue sections from control and experimental groups were processed on the same slide. Frozen sections (12 μm) were fixed in 4% paraformaldehyde in PBS for 10 min at room temperature and then stained with hematoxylin and eosin for light microscopy analysis. Images presented are representative of three independent experiments.

### Immunostaining

Staining FOXA2 (1:300, WRAB-FOXA2, Seven Hills Bioreagents), E-Cadherin (1:300, 3195s, Cell Signaling Technology), EGFR (1:100, 4267, Cell Signaling Technology), Ki67 (1:200, MA5-14520, Invitrogen) and CK8 (1:100, TROMA-1, Hybridoma Bank, Iowa) was performed using secondary antibodies conjugated with Alexa 488, or Alexa 594 (1:300, Jackson Immuno Research). Nuclear staining was performed using Hoechst 33342 (4 μg/ml, H1399, Thermo Scientific). Tissue sections from control and experimental groups were processed on the same slide for each experiment. Images presented are representative of three independent experiments.

### Whole-mount Immunostaining for 3D imaging

To reveal the tridimensional visualization of implantation sites, whole mount immunostaining with 3DISCO clearing was performed as previously described (18). Anti-E-Cad antibody (1:100, 3195s, Cell Signaling Technology) was used to stain the luminal epithelium. 3D images were acquired by a Nikon upright confocal microscope (Nikon A1R). To construct the 3D structure of the tissue, the surface tool in Imaris (Bitplane) was used.

### Fluorescence *in situ* hybridization

DIG-labeled probes were generated according to the manufacturer’s protocol (Roche). PFA-fixed frozen sections from control and experimental groups were hybridized with digoxigenin (DIG)-labeled cRNA probes. Frozen sections (12 μm) from each genotype and treatment group were processed on the same slide for each probe. Briefly, following fixation (in 4% PFA/PBS) and acetylation, slides were hybridized at 55°C with DIG-labeled *Lif* and *Msx1* probe. Anti-Dig-peroxidase was applied onto hybridized slides following washing and peroxide quenching. Color was developed by TSA (Tyramide signal amplification) Fluorescein according to the manufacturer’s instructions (PerkinElmer). Nuclear staining was performed using Hoechst 33342 (4 μg/ml, H1399, Thermo Scientific). Images presented are representative of three independent experiments.

### *In situ* hybridization using radioactive probes

*In situ* hybridization using radioactive (^35^S GTP) labelled *Lif* probes was performed as previously described (34). In brief, frozen sections (12 μm) were mounted onto poly-L-lysine-coated slides and fixed in cold 4% paraformaldehyde in PBS. The sections were prehybridized and hybridized at 45°C for 4 hours in 50% (Vol/Vol) formamide hybridization buffer containing ^35^S-labeled antisense RNA probes (Perkin Elmer). RNase A-resistant hybrids were detected by autoradiography. All sections were post-stained with H&E. Images presented are representative of three independent experiments.

### Progesterone (P_4_) and estradiol-17b (E_2_) Assays

Mouse blood samples were collected at 9:00 am on days 4 and 8 of pregnancy. Serum was separated by centrifugation and stored at –80°C until analysis. Serum hormonal levels in the serum were measured by P_4_ or E_2_ EIA kit (Cayman chemical) as previously described (14).

### Statistics analysis

Each experiment was repeated at least 3 times using independent samples. Data are shown as mean ± SEM. Statistical analyses were performed using a two-tailed Student’s t-test. A value of *P*<0.05 was considered statistically significant.

## Author contributions

M.M., J.Y., Y.S.K. and A.D. performed experiments. M.M., J.Y., X.S. and S.K.D. designed experiments. M.M., J.Y., Y.S.K, X.S. and S.K.D. analyzed data. M.M., X.S., and S.K.D wrote the manuscript.

## Author Information

The authors declare no competing interest.

## Acknowledgments

We thank Katie Gerhardt for her excellent editing of the manuscript. This work was supported in parts by NIH grants (HD103475 and HD068524 to SKD).

---

**Supplemental figure 1.**
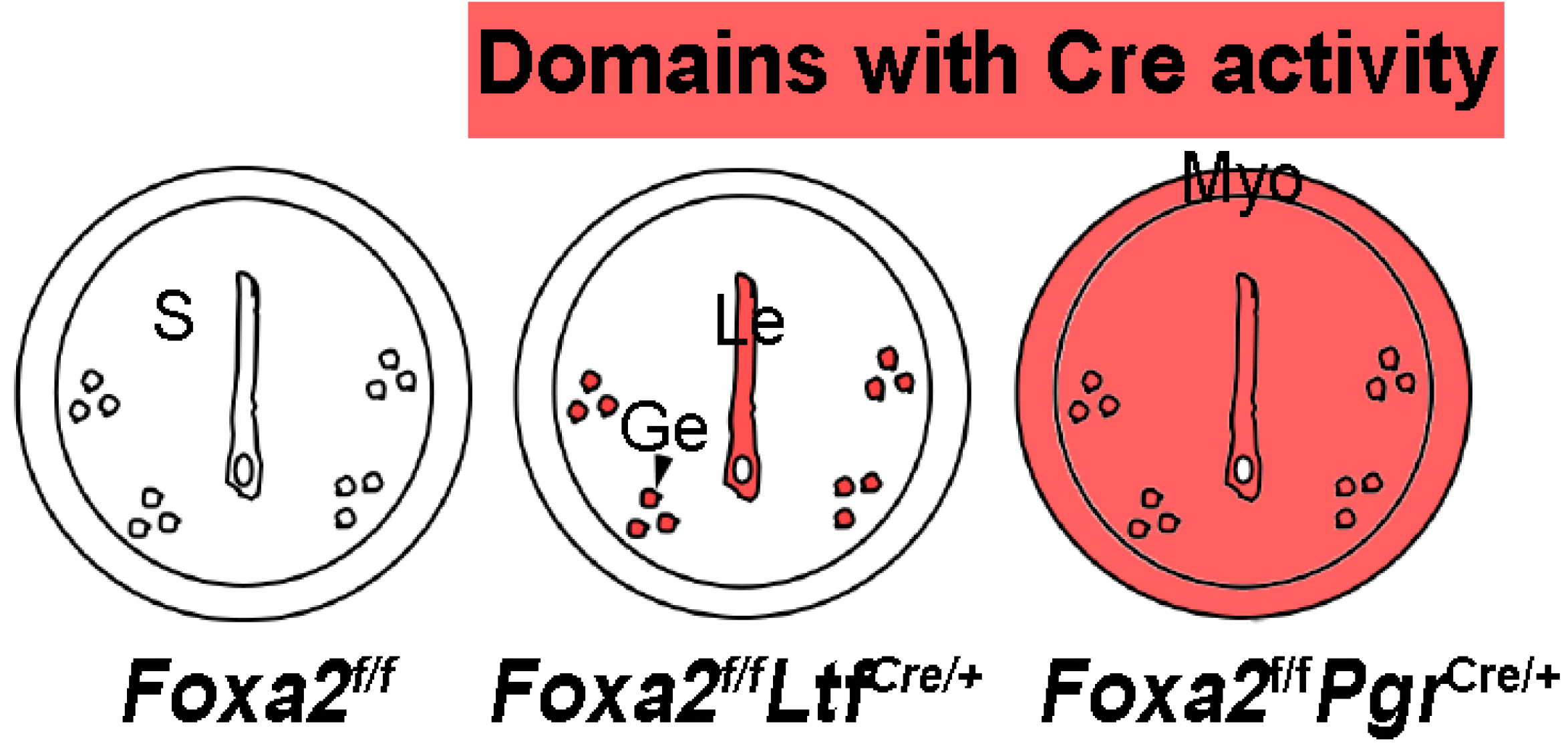
Scheme of Cre activity under *Ltf* and *Pgr* promoters. Domains with Cre activity are highlighted in red.

**Supplemental figure 2.**
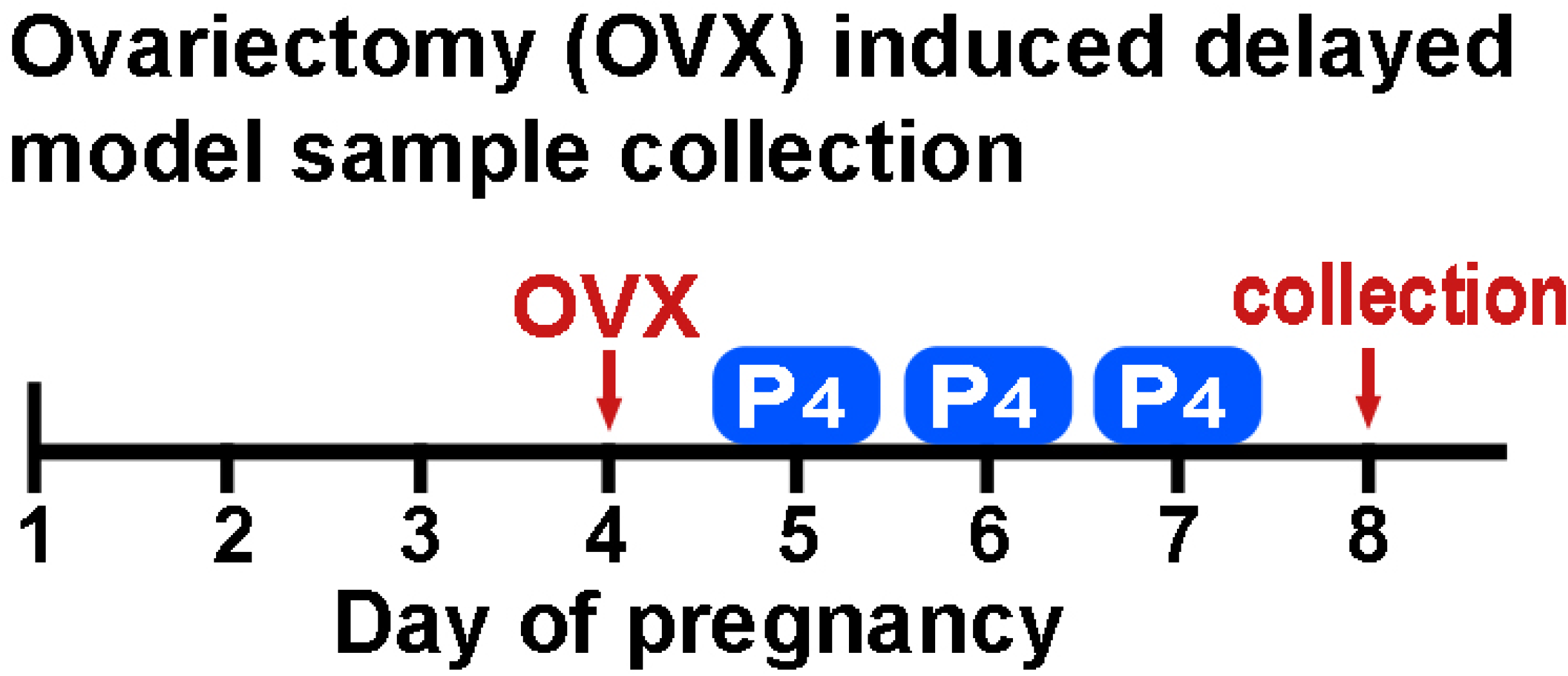
A schematic outline of sample collection from ovariectomy induced delayed model. P_4_, Progesterone (2mg) administration.

**Supplemental figure 3.**
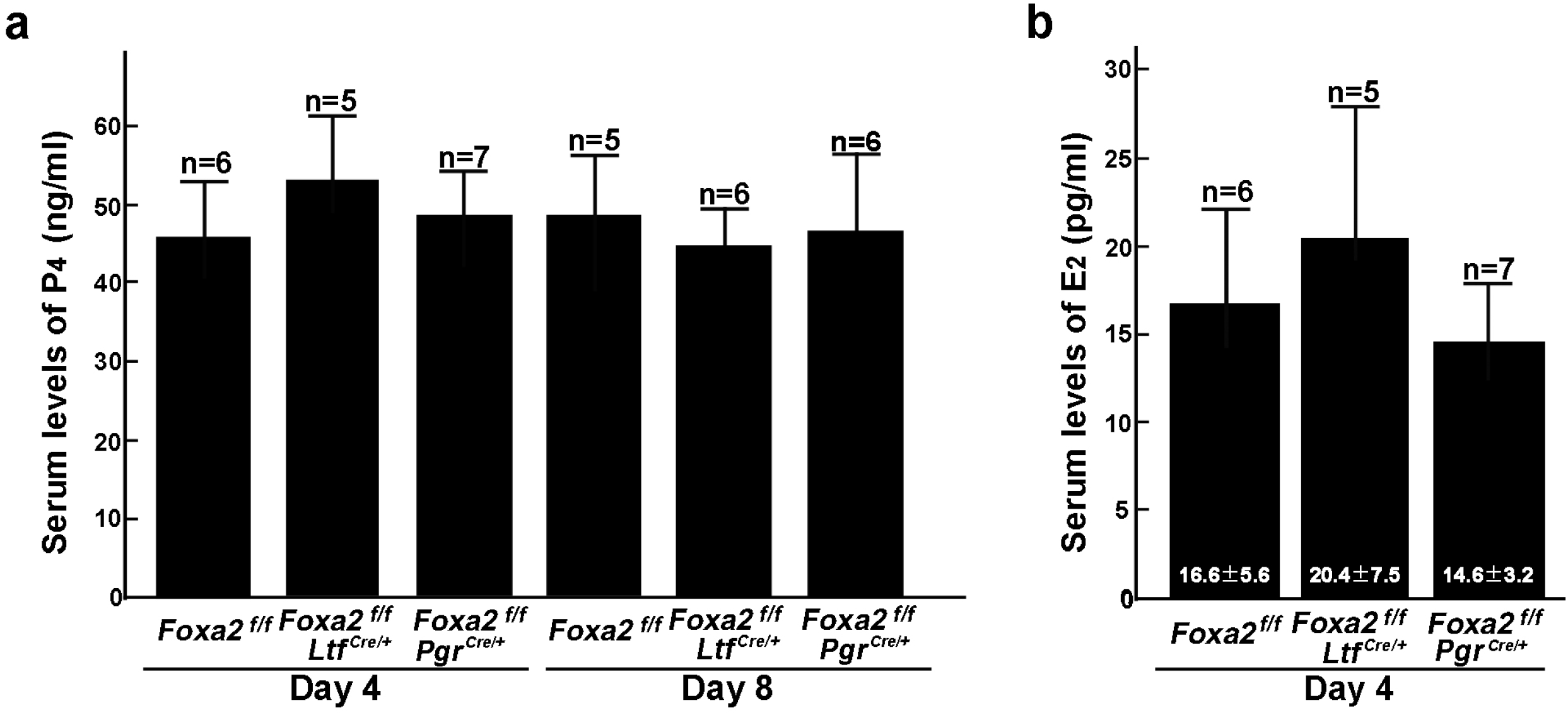
Serums levels of P_4_ and E_2_ in *Foxa2^f/f^*, *Foxa2^f/f^Ltf*^*Cre*+^ and *Foxa2^f/f^Pgr*^*Cre*+^ females on days 4 and 8 of pregnancy. Numbers of animals examined are presented on bars.

**Supplemental figure 4.**
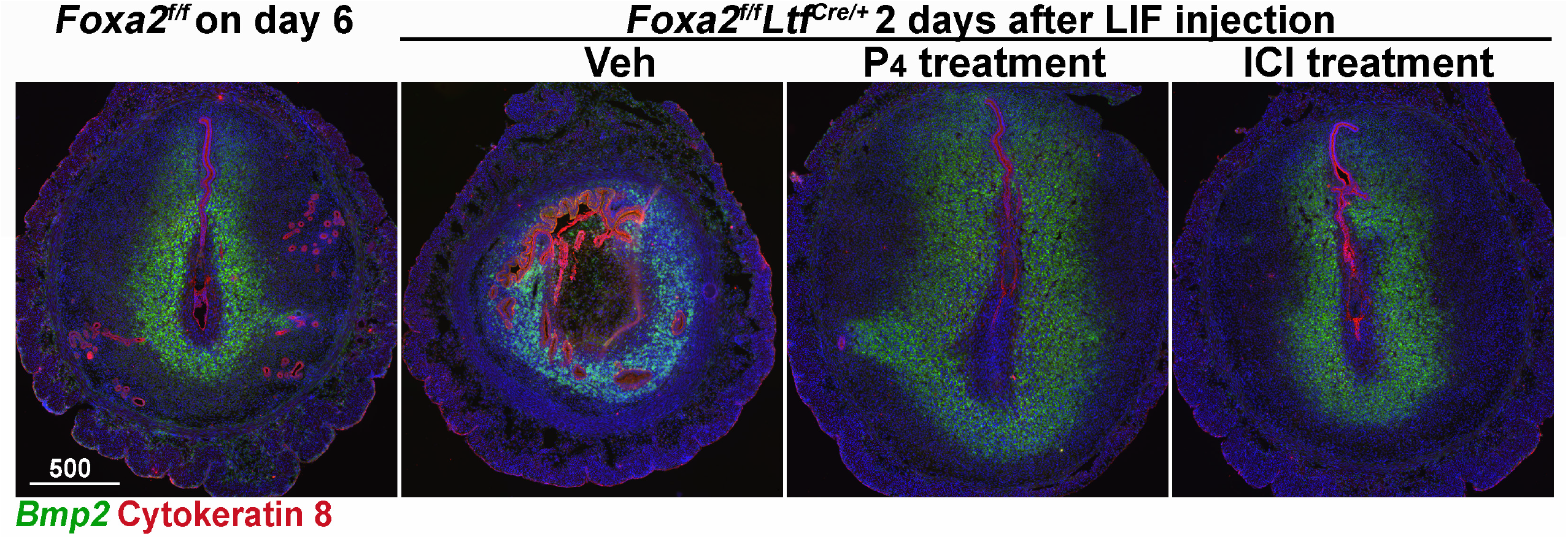
Fluorescence *in situ* hybridization of *Bmp2*. Implantation sites of *Foxa2^f/f^* females were collected on day 6 of pregnancy. Implantation sites of *Foxa2^f/f^Ltf*^*Cre*+^ females with different treatments were collected two days after LIF injections. Scale bar: 500 μm.

**Supplemental table 1.** Implantation sites in *Foxa2^f/f^*, *Foxa2^f/f^Ltf*^*Cre*+^ and *Foxa2^f/f^Pgr*^*Cre*+^ females on day 8 of pregnancy.

|  | No. of mice | No. of mice with IS (%) | No. of IS | No. of mice without IS (%) | No. of blastocysts recovered |
| --- | --- | --- | --- | --- | --- |
| ***Foxa2<sup>f/f</sup>*** | 6 | 6 (100%) | 11.5 ± 0.3 | 0 (0%) | N/A |
| ***Foxa2<sup>f/f</sup>Ltf<sup>Cre/+</sup>*** | 6 | 1 (16.7%) | 8 | 5 (83.3%) * | 5.4 ± 0.7 |
| ***Foxa2<sup>f/f</sup>Pgr<sup>Cre/+</sup>*** | 7 | 3 (42.9%) | 4.6 ± 1.2 | 4 (57.1%) * | 4.0 ± 0.3 |

\**P* < 0.05, Fisher's exact probability test.

